# Phenotypic Screening Coupled with AI-Driven Target Deconvolution Identifies α-Terthienyl as a Dual DPP-IV/HSD17β13 Modulator with Efficacy in a Mouse Model of MASLD

**DOI:** 10.64898/2025.12.12.693988

**Authors:** Kathryn M. Hoegeman, Jesse W. Wotring, Reid Fursmidt, Jedidiah Gaetz, Ehab M. Khalil, Douglas W. Selinger, Ilya Kovalenko, Tracey L. Schultz, Sean M. McCarty, Matthew J. O’Meara, Martin C. Clasby, Jonathan Z. Sexton

## Abstract

**Background:** Metabolic dysfunction-associated steatotic liver disease (MASLD) is a highly prevalent condition characterized by fat build-up in the liver and ranges from benign steatosis to progression to metabolic dysfunction-associated steatohepatitis (MASH), fibrosis, cirrhosis and end-stage liver disease including hepatocellular carcinoma, representing a significant cause of chronic liver disease globally.^1,2^ Current treatment options are limited, primarily relying on lifestyle modifications, highlighting an urgent need for novel therapeutic strategies.

**Methods:** A Cell Painting-style high-content screening phenotypic assay was employed using the PH5CH8 human hepatocyte cell line to identify small molecules capable of modulating induced hepatic steatosis. The Plex Research artificial intelligence (AI) platform was utilized for target deconvolution of the lead hit compound, α-terthienyl. In vivo efficacy was assessed in a diet-induced obesity (DIO) C57BL/6J mouse model of MASLD. Biochemical assays and molecular docking simulations were performed to validate predicted target interactions.

**Results:** Phenotypic screening identified 15 chemical probes/drugs that elicit dose-responsive inhibition of steatosis, and 16 that exacerbate steatosis, which could contribute to worsening of the disease clinically. α-terthienyl, a plant-derived natural product, was identified as a potent and non-toxic inhibitor of steatosis in PH5CH8 cells with an EC_50_ of 106 nM. In vivo, α-terthienyl administration to diet-induced obesity (60% fat diet) mice significantly reduced hepatic steatosis histologically, improved glucose tolerance, and favorably modulated serum biomarkers including ALT and AST. AI-driven analysis predicted dipeptidyl peptidase 4 (DPP-IV) and 17-beta hydroxysteroid dehydrogenase 13 (HSD17β13) as potential molecular targets of α-terthienyl. Biochemical inhibition of DPP-IV was observed and an oxidized α-terthienyl analog inhibited HSD17β13. Molecular docking supported these predictions, indicating binding to DPP-IV and HSD17β13.

**Conclusion:** This study demonstrates the successful application of phenotypic screening integrated with AI-driven target deconvolution to identify compounds and drugs that ameliorate or exacerbate hepatic steatosis. α-terthienyl was identified as a novel modulator of hepatic steatosis with in vivo efficacy in a MASLD model. Our findings suggest a dual-target mechanism involving DPP-IV and HSD17β13, potentially engaged by the parent compound and its metabolite, respectively, offering a promising polypharmacological approach for MASLD treatment.

## 1. Introduction

The metabolic syndrome epidemic describes a combination of interrelated disorders associated with obesity including hyperglycemia, insulin resistance, hypertension, and elevated plasma triglycerides. While type 2 diabetes mellitus (T2DM) is one of the most visible health consequences, the substantial impact on the liver is less appreciated. Normally, excess fatty acids accumulate as triglycerides in adipose tissue; however, ectopic triglyceride accumulation in non-adipose tissue can impair cellular function leading to lipotoxicity.^3^ Metabolic dysfunction-associated steatotic liver disease (MASLD) is characterized by excessive triglyceride deposits in the liver. The severe inflammatory stage of MASLD, metabolic dysfunction-associated steatohepatitis (MASH), occurs when this lipotoxicity leads to hepatocyte loss, decline of hepatic function, and fibrosis which in turn can cause cirrhosis, hepatocellular carcinoma (HCC) and acute liver failure.^3–5^ Strikingly, MASLD prevalence within the United States is estimated at around 38%, marking a 50% increase over the last three decades.^4^ Based on a meta-analysis of 86 MASLD epidemiological studies,^6^ the global prevalence of MASLD is slightly better but still quite serious at 25% with 40–76% of patients progressing to MASH and fibrosis that can result in end-stage liver disease and HCC.^7–9^ Beyond specific liver damage, MASLD is not only correlated with T2DM but can also contribute to diabetic hyperglycemia due to hepatic insulin resistance and correlates with T2DM as the majority of type 2 diabetics (50-60%) have some form of MASLD, suggesting that treatment of MASLD would positively impact T2DM.^10^ As of 2025, nearly 50% of the global population is overweight or obese and MASLD/MASH has become the leading cause of liver disease and driven dramatic increases in liver transplantation.^8,11–14^ While simple liver steatosis in the absence of fibrosis is considered to be a relatively benign condition, epidemiological studies suggest that the presence of fibrosis predicts both disease progression and liver-related complications over a subsequent 10-year period.^15,16^ Thus, one unifying therapeutic approach for MASLD, metabolic syndrome (MetS), and improving T2DM liver function may be decreasing ectopic triglyceride accumulation by stimulating fatty acid oxidation in the liver.

MASLD/MASH represents an impending global health epidemic, and the therapeutic landscape has significantly advanced in 2024–2025 with two distinct therapies now approved by the FDA, offering different mechanisms to treat MASH.^17^ The first, resmetirom (Rezdiffra), received accelerated approval in March 2024 for adults with noncirrhotic MASH and moderate to advanced liver fibrosis (stages F2 or F3). It acts as a liver-selective thyroid hormone receptor-β agonist that enhances fatty-acid metabolism and reduces fat accumulation.^18,19^ In August 2025, semaglutide (Wegovy) became the second drug to gain FDA approval for MASH. As a glucagon-like peptide-1 (GLP-1) receptor agonist, semaglutide works systemically by addressing the underlying metabolic syndrome. The apparent efficacy for GLP-1 receptor agonists in resolving steatohepatitis and improving fibrosis validates the approach of treating the metabolic insults that drive liver damage, marking a pivotal new treatment pathway for patients. Targeting associated comorbidities like T2DM or dyslipidemia may indirectly benefit liver health, but do not address the full complexity of MASLD/MASH pathogenesis. First, GLP-1 receptor agonists are not appropriate or tolerable for all individuals with MASLD. Second, although current therapies are approved for MASH with fibrosis, there remains a significant need for safe, liver-targeted agents that can prevent disease progression.

Developing therapeutics to treat multifactorial diseases like MASLD is challenging, as the underlying pathophysiology is not completely understood, involves multiple interacting pathways, and targeting single, specific proteins has yielded limited clinical success. Phenotypic drug discovery (PDD) offers an alternative paradigm to the dominant target-based drug discovery (TDD) approach. In PDD, compounds are screened for their ability to produce a desired biological effect, or phenotype, in a relevant cellular or organismal model, without *a priori* knowledge or bias regarding the specific molecular target.^20^ PDD is particularly well-suited for complex diseases^20^ as it evaluates compounds in a more physiologically relevant context, potentially identifying molecules that modulate complex pathways or possess polypharmacology beneficial for the disease state.^21^ Historically, PDD was responsible for the discovery of many first-in-class medicines.^21^ Driven by technological advancements in high-throughput imaging (e.g., high-content screening, "cell painting"), automated microscopy, and complex data analysis, PDD is experiencing a resurgence. However, a significant challenge remains: the subsequent identification of the molecular target(s) and mechanism of action (MoA) of the active "hit" compounds, a process known as target deconvolution.^21^ This step is crucial for lead optimization, understanding potential side effects, and regulatory approval. Moreover, because PDD relies on phenotypic readouts in model systems, it carries an elevated risk of identifying hits with spurious or non-specific phenotypic activity that may not translate in vivo, further underscoring the importance of rigorous mechanistic validation.

Given the urgent need for novel MASLD therapies and the potential of integrated drug discovery approaches, this study aimed to utilize high-content phenotypic screening in a human hepatocyte model to identify novel small-molecule modulators of hepatic steatosis. Additionally, this study focused on screening a “drug repurposing” collection of highly annotated compounds that are either FDA-approved, have been tested in human clinical studies, are in vivo active, or mechanistic chemical probes to both discover in vivo active compounds that modulate hepatic steatosis and to identify molecular pathways/targets that modulate the steatosis phenotype.

PDD screening of highly annotated chemical probes, FDA-approved drugs, and clinical candidate compounds can lead to the identification of efficacious compounds. A key challenge is identifying among all proteins and other macromolecules binding sites, which ones the compound directly interact with to cause the observed phenotypic effects. Clear target identification is important for translational drug discovery as it facilitates identifying the most appropriate patient population and developing biomarkers. Two complementary strategies for target deconvolution are bottom-up, where the physical interaction with each candidate target is considered and how that binding affects its specific molecular function; and top-down, where understanding of the signaling pathways and differential effects of the compound across different systems constrain the set of candidate targets. Integrating these approaches is challenging, as it often requires synthesis of diverse forms of evidence spread across the scholarly literature, public databases and knowledge repositories, and project-specific datasets. Excitingly, this data-integration bottleneck for target deconvolution in PDD is increasingly being addressed by the integration of artificial intelligence (AI) and machine learning (ML) techniques.^21^ In this study we have employed the Plex Research AI platform enabling drug-target enrichment analysis and to generate target hypotheses for hits.^22,23^

MASLD is a complex disease driven by multiple interconnected pathophysiological processes involving lipid metabolism, glucose regulation, inflammation, and fibrosis.^24^ Targeting a single pathway may be insufficient for effective treatment. Polypharmacology, the strategy of designing single molecules, known as multi-target-directed ligands (MTDLs) to intentionally engage multiple biological targets simultaneously, offers potential advantages over administering combinations of single-target drugs (polytherapy).^25^ Potential benefits include synergistic or additive efficacy, improved side effect profiles via lower effective doses for each target, reduced risk of drug-drug interactions, simplified dosing regimens leading to better patient compliance, and potentially lower development costs. This approach has shown considerable success recently in metabolic diseases, exemplified by dual and triple incretin receptor agonists for T2DM and obesity.^26^

Given the need for novel MASLD therapies and the potential of integrated drug discovery approaches, this study aimed to use high-content phenotypic screening in a human hepatocyte model to identify novel small-molecule modulators of hepatic steatosis. Additionally, this study focused on screening a “drug repurposing” collection of highly annotated compounds that are either FDA-approved, have been tested in human clinical studies, are in vivo active, or mechanistic chemical probes to both discover in vivo active compounds that modulate hepatic steatosis and to identify molecular pathways/targets that modulate the steatosis phenotype. Subsequently, the objective was to employ the Plex Research AI platform, which combines centrality algorithms and chemical similarity searching with leveraging publicly available data, knowledge graphs and integrated LLMs to find convergence between compound structures and diverse evidence types from large, publicly available datasets to aid in mechanistic interpretation and target hypothesis generation. The Plex AI platform guided target deconvolution and drug-target/pathway analysis of the in vivo-active hit compound, α-terthienyl, to predict its molecular mechanism. Finally, the study sought to validate the in vivo efficacy of α-terthienyl and provide biochemical and in silico support for its proposed AI-predicted targets and mechanism of action.

## 2. Materials and Methods

### 2.1 Cell Culture and Reagents

The PH5CH8 human hepatocyte cell line, derived from primary human hepatocytes immortalized via SV40 T antigen expression, was used for *in vitro* studies.^27^ Cells were maintained in Dulbecco’s Modified Eagle Medium (DMEM, Thermo Fisher Catalog #11995965) supplemented with 10% fetal bovine serum (FBS), 1% penicillin-streptomycin under standard cell culture conditions (37°C, 5% CO2). Cells were continuously evaluated for mycoplasma contamination, and the results were always negative. Cell passage numbers were monitored, and new cell stocks were thawed for follow-up experiments including concentration-response experiments.

### 2.2 Compounds

The library used for screening was the Selleckchem Bioactive Compound Library-II (Catalog No. L1700-II), containing 5,309 compounds dissolved in dimethyl sulfoxide (DMSO) at 10 mM. The library included FDA approved compounds, clinical candidates, and other highly annotated bioactive compounds. The library was reformatted from 96-well plates to 384-well plates and diluted to 2 mM prior to screening. Compounds identified as hits from the primary screen were repurchased from MolPort and solubilized at 10 mM in DMSO. All compounds were stored at -80°C and were kept out of the light. For the primary screen, compounds were added using a 50 nL pin tool at the University of Michigan Center for Chemical Genomics and were screened at a final concentration of 2 μM in cells. For concentration response follow up experiments, compounds were added using an HPD300e digital compound dispenser across a range of concentrations.

### 2.3 Poly-D-Lysine coating

Perkin Elmer Cell Carrier Ultra 384-well microplates (6057300, Perkin Elmer) were coated with Poly-D-Lysine (PDL, Santa-Cruz Cat# sc-136156) prior to cell plating to enhance cell attachment and reduce cell-detachment after fixation. 15 μL/well of 1 mg/mL PDL, solubilized in molecular grade biology water, was added to plates and allowed to incubate at 37°C and 5% CO_2_ for 1 hour. After incubation, plates were washed two times with molecular grade biology water and then allowed to air dry in a cell culture hood with the lid off prior to use.

### 2.4 Preparation of sodium oleate stock

15 mL aliquots of 100 mM sodium oleate stocks were generated by adding 457 mg sodium oleate (Sigma Aldrich Cat#03880) and 25 μL 1N NaOH to 15 mL ddH_2_O at warming to 37°C. 100 mM stocks were filtered using 0.22 μm filters and frozen at -20°C for storage prior to use. 40 mL aliquots of 10 mM sodium oleate working stock, loaded onto bovine serum albumin (BSA), was created by adding 4 mL of thawed 100 mM sodium oleate stocks dissolved in 13.4 mL BSA solution (30% BSA in DPBS, sterile-filtered, BioXtra, Sigma Aldrich A9576) and 22.6 mL serum free DMEM and warming to 37°C. These BSA complexed 10 mM oleate working stocks were what was directly added to cells for lipid loading in the high-content bioassay for hepatic steatosis.

### 2.5 High-Content Steatosis Screening Assay

To PDL coated plates, 20 μL of complete media was added to each well, and PH5CH8 cells were seeded on top at an optimized density of 1,750 cells/well in 25 μL/well of additional volume. Following seeding, cells were lipid loaded with 5 μL/well of 10 mM sodium oleate working stock to produce final concentration of 1 mM sodium oleate in each loaded well. 32 loaded control wells (negative control, NC) and 32 unloaded control wells (positive control, PC) were included on each plate. Unloaded controls were normalized to a constant assay volume of 50 μL/well with 5 μL/well additions of complete media. After loading, cells were allowed to incubate overnight at 37°C and 5% CO_2_. The next day, compounds were dispensed to wells at a final concentration of 2 µM and all wells were normalized to a constant 0.1% DMSO concentration. After 72 hours of compound treatment, the ER stain ER-blue (Thermo Fisher Cat# H34477) was added at 1:10,000 to live cells for 30 minutes in the incubator. Then, cells were fixed using 50 μL/well of 4% paraformaldehyde for 20 minutes, permeabilized with 0.3% Triton-X100 for 15 minutes, and rinsed 2X with PBS prior to staining with the optimized cell dye set containing 200 nM MitoView Green (Biotium, Cat# 70054), 10µg/mL Hoechst 33342 (Sigma Aldrich Catalog # H21492), 1:50,000 CellMask orange (Thermo Fisher Cat# H32713) and 1:2,000 HCS LipidTOX deep red neutral lipid stain (Thermo Fisher Cat# H34477) for 30 minutes at room temperature. After staining, cells were rinsed 3X with PBS, then left in a final volume of 50 μL/well PBS for imaging.

### 2.6 Fluorescence Imaging and Image Processing

Stained plates were imaged using either the Yokogawa Cell Voyager 8000 or a Thermo Fisher CX5 high-content microscope at 20x magnification. Excitation power and exposure times were optimized to achieve optimal signal-to-background ratio and images were collected at a single Z-plane as determined by autofocus on the DNA channel. A total of 6 fields were captured for each well for analysis.

An automated image analysis pipeline was employed to segment cells and nuclei and extract many quantitative features describing cell morphology, intensity, and texture, including metrics specifically quantifying lipid droplet content (e.g., total lipid droplet count, area per cell, lipid droplet intensity). Cellpose (version 3.1.1)^28^ was used for nuclear segmentation, and CellProfiler (version 4.2.6)^29^ was used for feature extraction. Cellular features/objects identified included nuclei, mitochondria, endoplasmic reticulum, nucleoli (visible in the HCS CellMask Orange channel), cytoplasm/cytoskeleton, and neutral lipid droplets. Each of the identified sub-cellular objects, including individual lipid droplets, were related back to a parent cell using the “relate objects” module. Tabulated measurements included texture, area/shape morphometrics, intensity, radial distribution, etc. for each object. A total of approximately 500 unique features were extracted for each identified cell (feature vector) and mean/median results were summarized to the treatment-well level for machine learning scoring.

### 2.7 Primary Screen Scoring Using Machine Learning

A random forest regression model (ML score) was trained using XGBoost with the 500 cellular feature vectors on positive and negative control wells, representing 100% or 0% effect, respectively.^30^ Data preprocessing included centering/scaling of the feature columns, batch correction using the HarmonyPy package, and highly correlated features above 95% correlation were pruned and low variance features (<2%) were also pruned.

The model was trained using 80% of the data and model performance was evaluated on the remainder 20% of the withheld test data (to control for over-fitting). The sensitivity/specificity on the test data set was greater than 99%. Hits were identified based on statistically significant changes in the ML score compared to vehicle controls (e.g., Z-score < +3 for inhibitors, > -3 for exacerbators). Additionally, the percent viability for each compound was calculated with the following formula: Percent Viability = (Cell Count / Average DMSO Control Cell Count) × 100%.

### 2.8 Dose-Response Confirmation

Dose-response experiments were conducted for hits to determine potency (EC_50_) and cytotoxicity (CC_50_) via 4-parameter logistic regression using GraphPad Prism. A 10-point, 2-fold concentration series (N=1 replicate per condition) was tested for each compound ranging from 10 µM–40 nM. Imaging and analysis were performed as in the primary screen. Compounds with robust dose-responsiveness were repeated in triplicate to generate final dose-response curves.

### 2.9 Biochemical Assays

DPP-IV activity was measured using the DPP-IV-Glo™ Protease Assay (Promega, Cat. No. G8350) following the manufacturer’s instructions. The assay employs the proluminescent substrate Gly-Pro-aminoluciferin, which releases aminoluciferin upon DPP-IV cleavage, generating a luminescent signal proportional to enzymatic activity. HSD17B13 inhibitory activity was evaluated using the NAD(P)H-Glo™ Detection System (Promega, Cat. No. G9061) according to the manufacturer’s instructions. This bioluminescent assay quantifies reduced nicotinamide cofactors (NADH and NADPH) to assess enzyme activity and inhibition.

### 2.10 Drug Target Enrichment and Deconvolution (Plex Research Platform)

AI-driven target deconvolution was performed using the Plex Research platform^22^ that integrates multi-modal data, including compound chemical structures, quantitative phenotypic fingerprints, and extensive biological knowledge graphs from publicly available data. Central to this process is the generation of "focal graphs," knowledge subgraphs focused on the query compound and its structurally related entities, which are analyzed using centrality algorithms (like PageRank) to identify potential protein targets based on convergence of diverse evidence within the graph. This approach yields a transparent, ranked list of target hypotheses with associated confidence scores, enabling rapid target prioritization and enrichment analysis. Target enrichment analysis was performed for a selection of 15 steatosis inhibitors and 16 exacerbators hits from the phenotypic screen for prioritization with further empirical assessment of molecular targets/pathways that regulate hepatic steatosis in this in vitro model.

### 2.11 Molecular Docking Simulations

Molecular docking simulations were conducted to assess the plausibility of interactions between α-terthienyl with the predicted targets, DPP-IV and HSD17β13. Docking was performed using AutoDock Vina. Crystal structures of human DPP-IV (PDB ID: 1NU6) and human HSD17β13 (PDB ID: 8G9V) were obtained from the Protein Data Bank. Protein structures were prepared by removing water molecules and existing ligands, adding hydrogen atoms, and assigning appropriate protonation states and charges. The binding site was defined based on the location of co-crystallized ligands or predicted active site cavities. The 3D structure of α-terthienyl was generated and energy-minimized using standard chemistry software. The ligands were prepared for docking by assigning appropriate atom types and charges. Flexible ligand-rigid receptor docking was performed.^31^ Multiple docking runs were initiated using a genetic algorithm or similar search method to explore conformational space within the defined binding site. The resulting poses were ranked using the software’s scoring function (Vina score), which estimates the binding free energy or affinity.^32^

### 2.12 Animal Studies

All animal procedures were approved by the University of Michigan’s Institutional Animal Care and Use Committee and conducted in accordance with established ethical guidelines. Male C57BL/6J mice (The Jackson Laboratory, Stock No: 000664) were used because of their predisposition to hepatic steatosis and pre-diabetes.^33,34^ At 6 weeks of age, mice were randomly assigned to either a control diet (Research Diets D12450B, 10 kcal% fat) or a high-fat diet (HFD; e.g., Research Diets D12492, 60 kcal% fat) to induce obesity and MASLD features. Mice were maintained on these diets for 16 weeks to establish the DIO-MASLD phenotype, characterized by obesity, glucose intolerance, insulin resistance, and hepatic steatosis.^34^ Following the diet induction period, DIO mice were randomly assigned to receive either vehicle control or α-terthienyl at 1, 5, and 20 mg/kg body weight via intraperitoneal injections daily for 14 days. Body weight and food intake were monitored regularly.

An intraperitoneal glucose tolerance test (GTT) was performed both before and after the treatment period. Mice were fasted for 6 hours, followed by intraperitoneal injection of D-glucose at 2 g/kg body weight. Blood glucose levels were measured from tail vein blood at baseline (0 min) and at 15-, 30-, 60-, 90-, and 120-minutes post-injection using Contour Next blood test strips and glucometer. At the study endpoint, mice were euthanized via exsanguination. Blood was collected via cardiac puncture for serum biomarker analysis, and livers were excised, weighed, and processed for histology and other analyses.

### 2.13 Histological Analysis

Liver tissue samples were fixed in 10% neutral buffered formalin, processed routinely, and embedded in paraffin. 5 µm thick sections were cut and stained with Hematoxylin and Eosin (H&E). Stained slides were digitized using an Aperio whole-slide scanner. Histopathological evaluation was graded based on the percentage of parenchymal area affected by macrovesicular and/or microvesicular steatosis (Score 0: <5%; Score 1: 5–33%; Score 2: 34–66%; Score 3: >66%).

### 2.14 Statistical Analysis

Data are presented as mean ± standard error of the mean (SEM) or standard deviation (SD) as indicated. Statistical comparisons between two groups were performed using an unpaired Student’s t-test. Comparisons among multiple groups were made using one-way analysis of variance (ANOVA) followed by appropriate post-hoc tests (Dunnett’s) for multiple comparisons. A p-value of < 0.05 was considered statistically significant.

Statistical analyses were performed using GraphPad Prism software (Version 10.2.3).

## 3. Results

### 3.1 High-Content Phenotypic Screening Identifies Modulators of Hepatic Steatosis

To identify compounds modulating hepatocyte lipid accumulation, a high-content phenotypic screening assay was developed using the PH5CH8 human hepatocyte cell line. To induce a steatotic phenotype, cells were exposed to 1 mM of the unsaturated fatty acid sodium oleate (**Figure 1A**) using fatty acid-free BSA as a carrier. PH5CH8 cells naturally have a low neutral lipid content, but when exposed to 1 mM sodium oleate, they accumulate roughly 30–40 lipid droplets per cell (**Figure 1B**). This assay was designed to accommodate one day of lipid loading/cell attachment and three days of drug treatment for a total assay window of 96 hours. At the end of the drug exposure period, cells were fixed and stained with a multiplexed fluorescent dye set including markers for cell nuclei (Hoechst 33342), endoplasmic reticulum (ER Tracker blue/white), cytoplasm (HCS CellMask orange), mitochondria (MitoView green), neutral lipid droplets (HCS Lipidtox Deep Red), and transmitted light to capture a wide range of cellular phenotypic features relevant to MASLD and overall cell health to enable selection of selective steatosis inhibitors/exacerbators independent of general cytotoxic effects. **Figure 1C** shows representative images of unloaded and loaded control cells in each fluorescent channel with the most notable difference in neutral lipid droplets upon stimulation with oleate.

**Figure 1.**
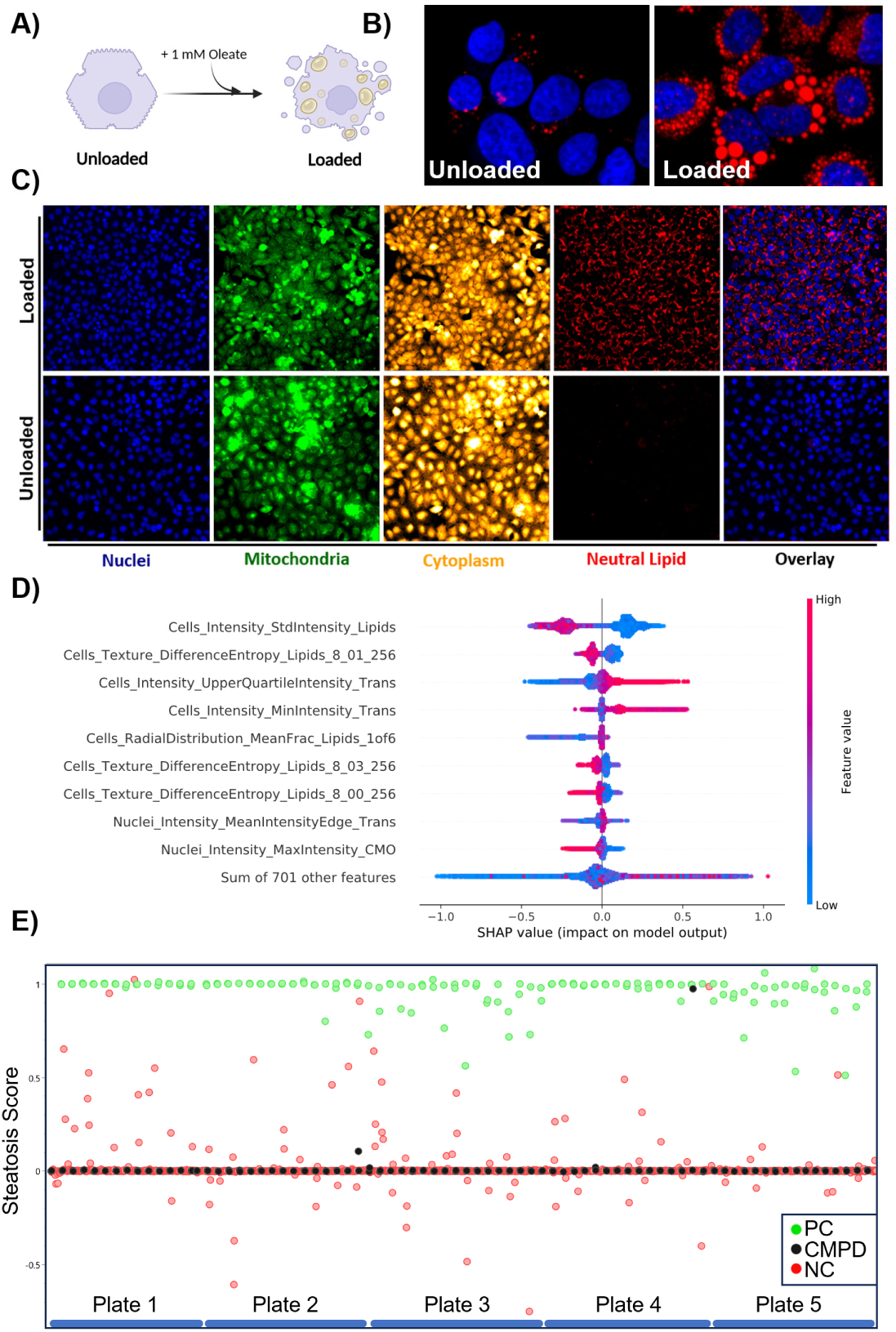
Cell painting-style assay for MASLD. A) Lipid loading - 1 mM of sodium oleate was used to lipid load PH5CH8 cells. B) Fluorescence images of unloaded (left) and loaded (right) PH5CH8 cells (nuclei=blue, lipids-red). C) Representative 20X images are shown for both unloaded and loaded fields of cell nuclei (blue), mitochondria (green), cytoplasm (orange), and neutral lipids (red). D) SHAP feature importance for the XGBoost steatosis scoring metric. E) Scatterplot for 5 plates showing negative controls, positive controls and tested compounds (CMPD).

A highly annotated drug library consisting of 5,309 FDA-approved drugs, chemical probes, and clinical candidate compounds with known targets/mechanisms of action was screened at 2 µM in 17 384-well plates and compounds that significantly inhibited (**Figure 1E** positive score) or exacerbated (**Figure 1E** negative score) the steatotic phenotype were selected for dose-response confirmation. To enhance sensitivity/specificity and robustness in primary screening for hit selection, an XGBoost machine learning model was trained on NC and PC conditions in lieu of selecting a single feature (e.g. lipid droplet count/size). Cell level features were calculated by using Cellpose for segmentation and CellProfiler. Cell level features were then centered/scaled. To facilitate interpretation, features with low variance (<5%) and or highly correlated with other features (>95%) were pruned yielding ∼500 features for approximately 1 million cells per plate. To control for over-fitting, the data was split (80%/20%) into training and test sets. HarmonyPy was applied to the data to correct for plate-based batch effects before model fitting. Model performance was then evaluated on the test set and an aggregated steatosis score per-well was tabulated. A SHAP (SHapley Additive exPlanations) analysis showing lipid droplet features, transmitted light intensity features, and CellMask features contribute significantly to the steatosis score (**Figure 1D**). After model training on controls, each well was scored and **Figure 1E** shows a scatter plot of five plates from the primary screen with positive controls labeled in green, negative controls labeled in red, and compounds labeled in black. Each plate was assessed for assay quality by comparing the lipid loaded, DMSO vehicle-treated negative controls (NC) to the non-lipid loaded, DMSO vehicle-treated positive controls (PC) and plates consistently exhibited a robust Z’ score > 0.5 and a total of 113 hits were identified in this screen that met the significance threshold. **Table 1** shows an overview of the selected dose-responsive steatosis inhibitors with α-terthienyl emerging as a compound of interest, demonstrating a potent, dose-dependent reduction in intracellular lipid content (dose-response curves shown in **Supplementary Figure 1**). Several compounds were also identified that significantly increased lipid accumulation compared to vehicle controls, providing a set of bidirectional modulators for subsequent analyses.

**Table 1:** Summary of steatosis inhibitors with their respective compound class and MASLD association.

| Compound | EC<sub>50</sub> (μM) | CC<sub>50</sub> (μM) | Compound Class | MASLD Association |
| --- | --- | --- | --- | --- |
| α-terthienyl | 0.106 | >10 | Natural Product<sup>35</sup> | No |
| CC-115 | 13.17 | >10 | mTOR Inhibitor<sup>36</sup> | No |
| CCG-1423 | 5.21 | >10 | Rho-A Inhibitor<sup>37</sup> | No |
| Digoxin | 4.57 | 0.052 | Cardiac Glycoside<sup>38</sup> | Yes |
| Dihydrocoumarin | 44.34 | >10 | Chromanone<sup>39</sup> | No |
| Domiphen Bromide | 9.94 | 4.90 | Antiseptic<sup>40</sup> | No |
| iCRT3 | 69.34 | >10 | Wnt Inhibitor<sup>41</sup> | No |
| LF3 | 84.50 | >10 | Wnt Inhibitor<sup>42</sup> | No |
| Ouabain | 4.849 | 0.023 | Cardiac Glycoside<sup>43</sup> | No |
| Pentoxifylline | 5.83 | >10 | Vasodilator<sup>44</sup> | Yes |
| Periplocin | 0.096 | 0.039 | Cardiac Glycoside<sup>45</sup> | No |
| Phthalyl Sulfacetamide | 621.7 | >10 | Antibiotic<sup>46</sup> | No |
| Rhodamine B | 2.81 | >10 | Dye<sup>47</sup> | No |
| Sorbic Acid | 14.85 | >10 | Preservative<sup>48</sup> | Yes |
| THZ1 | 0.969 | 0.034 | CDK7 Inhibitor<sup>49</sup> | No |

### 3.2 Target Enrichment Analysis Implicates Multiple Pathways in Hepatic Lipid Regulation

Using the known targets of all validated hit compounds (15 inhibitors and 16 exacerbators), we used the Plex Research platform to predict enriched biological pathways. This revealed significant enrichment for 15 biological pathways and target classes known to be involved in liver metabolism and MASLD pathogenesis (**Table 2**). Enriched terms included pathways related to lipid metabolism (e.g., fatty acid synthesis, fatty acid oxidation, cholesterol biosynthesis), glucose metabolism/insulin signaling, inflammatory signaling, and targets associated with nuclear receptors (e.g., PPARs) and metabolic enzymes. **Table 2** illustrates that several pathways, including Wnt/β-catenin signaling and autophagy/lipophagy, can either increase or decrease steatosis depending on the direction of perturbation. In contrast, inhibition of mTOR signaling consistently decreased steatosis, whereas more generalized disruption of cell-cycle regulation and transcriptional control reliably worsened the phenotype.

**Table 2.**
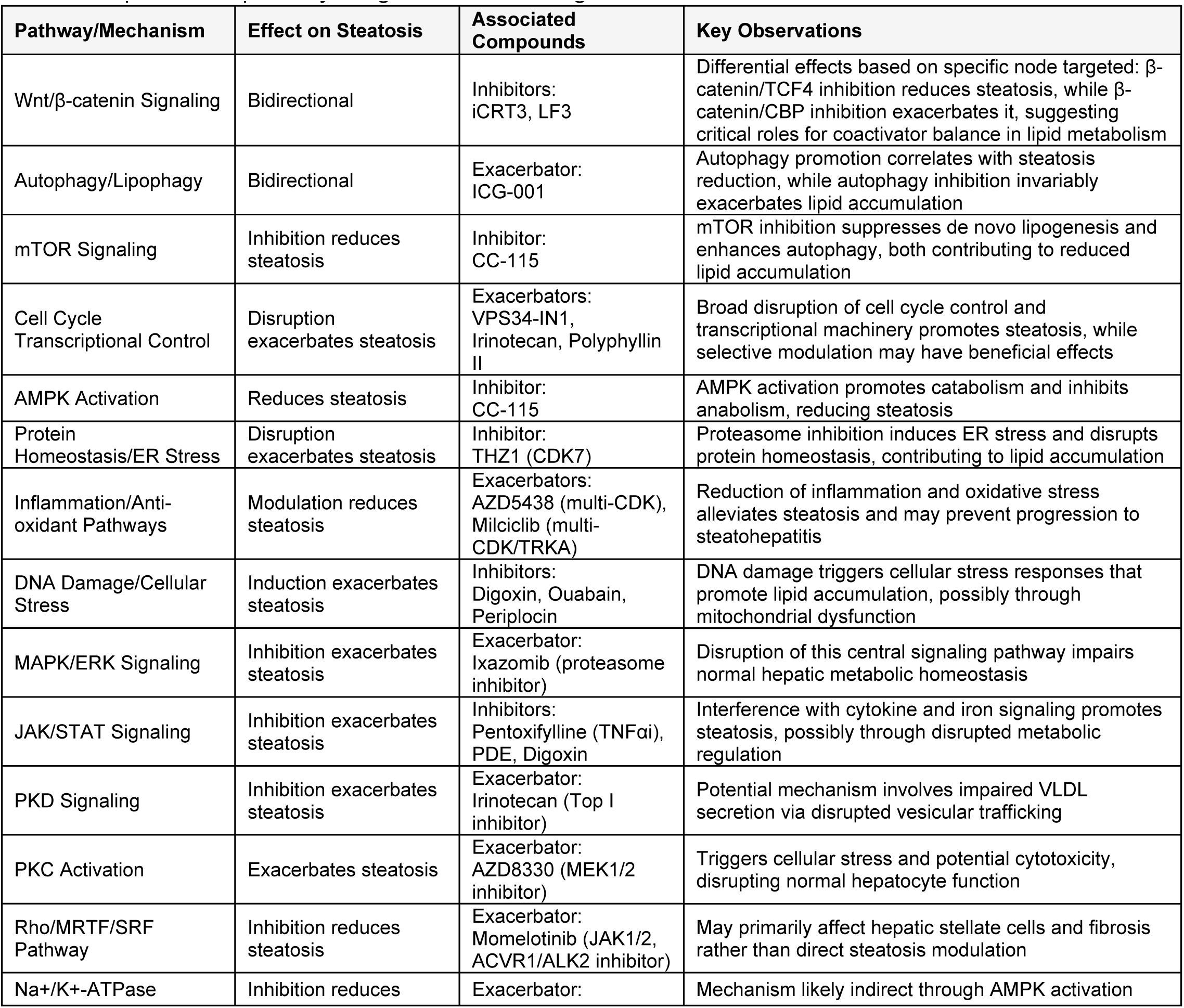
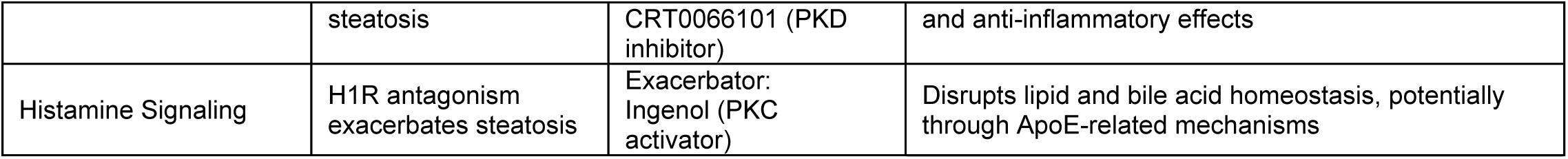
Top enriched pathways/targets from screening hits.

Analysis of this high-content screening data reveals distinct mechanistic patterns between compounds that inhibit versus exacerbate hepatic steatosis. Steatosis inhibitors primarily target pathways that restore metabolic homeostasis, including AMPK activation (cardiac glycosides like Digoxin/Ouabain), mTOR inhibition (CC-115), selective Wnt/β-catenin modulation (iCRT3/LF3 targeting β-catenin/TCF4), and anti-inflammatory mechanisms (Pentoxifylline). These compounds generally act by countering pathological signaling or enhancing protective pathways, targeting proteins or pathways whose dysregulation contributes to the metabolic imbalances characteristic of MASLD/MASH. For instance, mTOR inhibition suppresses de novo lipogenesis while promoting autophagy-mediated lipid clearance,^50,51^ while anti-inflammatory effects may mitigate the progression from simple steatosis to steatohepatitis.^52,53^

Conversely, steatosis exacerbators (dose-response curves shown in **Supplementary Figure 2**) predominantly disrupt essential cellular homeostatic processes, with notable convergence on autophagy inhibition (VPS34-IN1), proteasome inhibition (Ixazomib), DNA damage induction (Irinotecan), and cell cycle disruption (AZD5438, Milciclib, MK-5108). These compounds frequently target proteins or pathways essential for normal cellular function, maintenance, and stress responses, suggesting that compromised cellular homeostasis and integrity contribute significantly to pathological lipid accumulation. The exacerbation of steatosis through disruption of autophagy is particularly noteworthy given the established role of lipophagy in clearing hepatic lipid droplets, while proteasome inhibition triggers endoplasmic reticulum stress, a recognized contributor to hepatic steatosis development.^54–58^ Disruptions to these cellular homeostatic mechanisms represent critical pathogenic pathways in drug-induced liver injury (DILI), as compromised autophagy, proteasome function, and DNA repair capacity collectively impair the hepatocyte’s ability to maintain cellular homeostasis under xenobiotic stress. The connection of these pathways to steatosis exacerbation is significant for DILI pathogenesis where these mechanisms are primary initiating events in the hepatotoxicity cascade.^59,60^

A striking finding is the differential impact of targeting distinct nodes within the same pathway, exemplified by the Wnt/β-catenin signaling axis. While inhibitors of β-catenin/TCF4 interaction (iCRT3, LF3) reduce steatosis, the β-catenin/CBP inhibitor ICG-001 exacerbates lipid accumulation, suggesting critical roles for coactivator balance (CBP/p300) in hepatic lipid metabolism regulation. This mechanistic divergence underscores the complexity of transcriptional regulation in lipid homeostasis and highlights the potential therapeutic value of selective pathway modulation. Furthermore, the observation that different kinase inhibitors yield opposing effects on steatosis—mTOR inhibition being anti-steatotic while inhibition of MEK, multiple CDKs, Aurora kinases, JAKs, PKD, and Vps34 exacerbates lipid accumulation, which emphasizes the context-dependent roles of signaling networks in liver metabolism.

The enrichment of cellular stress response modulators among exacerbators highlights the balance between metabolic adaptation and cellular dysfunction in hepatic steatosis pathophysiology. The data suggest that while targeted inhibition of specific pathways can alleviate steatosis, broad disruption of essential cellular processes invariably promotes lipid accumulation, potentially through triggering stress responses that manifest as dysregulated lipid metabolism. These mechanistic insights deepen our understanding of the molecular underpinnings of hepatic steatosis but also identify promising therapeutic targets and approaches for MASLD/MASH intervention, particularly those focused on enhancing catabolic pathways, suppressing anabolic pathways, promoting cellular quality control, and reducing inflammation/oxidative stress in a highly targeted manner.

### 3.3 α-Terthienyl Ameliorates Steatosis and Improves Metabolic Parameters in a DIO Mouse Model of MASLD

The most potent and non-toxic hit from this screening campaign was α-terthienyl, which had an EC_50_ value of 106 nM and a 50% cytotoxic concentration (CC_50_) greater than 15 μM (**Table 1**). α-terthienyl, also known as 2,2’:5’,2”-terthiophene, is a naturally occurring compound found in African marigold roots with a simple structure composed of three consecutive thiophene rings (**Figure 2A**).^61,62^ To evaluate its anti-steatosis activity, we ran α-terthienyl in an additional concentration-response experiment, this time as a 20-point 1.5-fold dilution series from a maximum concentration of 15 μM (N=3 replicates per condition). Machine learning-based steatosis scoring was used to evaluate efficacy to discriminate positive and negative control conditions (**Figure 2B**), and high efficacy/potency was confirmed The EC_50_ was determined to be similar at 130 nM (**Figure 2A**), while the CC_50_ was higher than 15 μM (data not shown).

**Figure 2.**
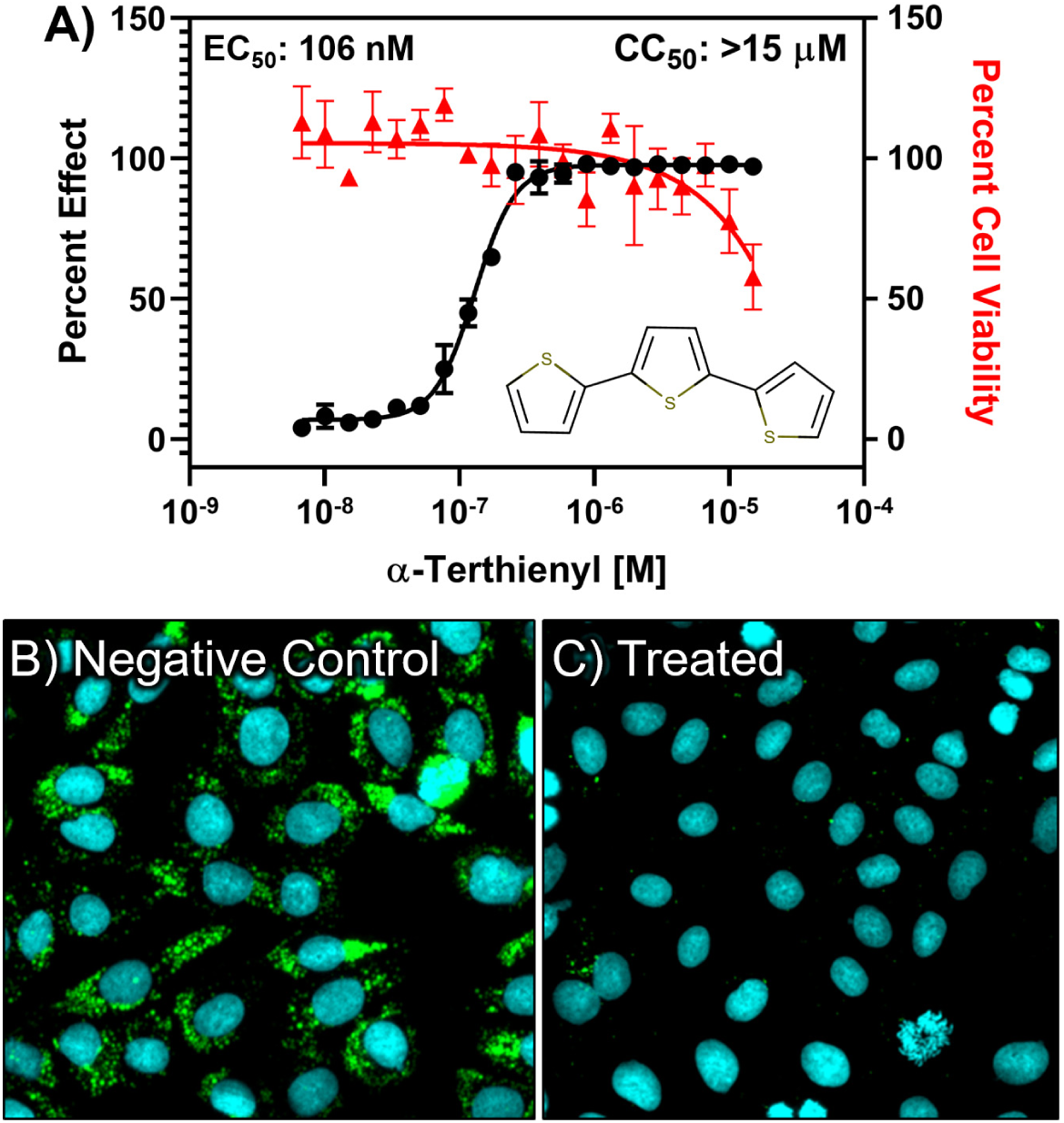
*In vitro* steatosis (black) and viability (red) dose response curves for α-terthienyl and representative images shown for B) vehicle control and C) α-terthienyl treated cells.

To evaluate the in vivo efficacy of α-terthienyl, the C57BL/6J mouse model of diet-induced obesity (DIO) and MASLD was used.^63^ 25-week old mice were purchased from Jax labs and were maintained on a 60% HFD for an additional 6 weeks. α-terthienyl had been previously evaluated for safety in vivo, where it was shown to have a 50% lethal dose (LD_50_) in rats of 110 mg/kg by intraperitoneal (IP) injection.^64^ We performed a 14-day study with daily IP injections of 5 mg/kg α-terthienyl (N=5) or vehicle (PBS containing 5% Cremophor EL, 20% PEG-300, and 20% DMSO) (N=7). Following the treatment period, IP blood glucose tolerance was evaluated and then animals were euthanized, livers were excised, weighed and processed for histological analysis (**Figure 3**). Remarkably, α-terthienyl had a strong, statistically significant effect in vivo. Treated animals had significantly smaller livers (**Figure 3D**), suggesting resolution of steatosis, which was further confirmed by liver histology.

**Figure 3.**
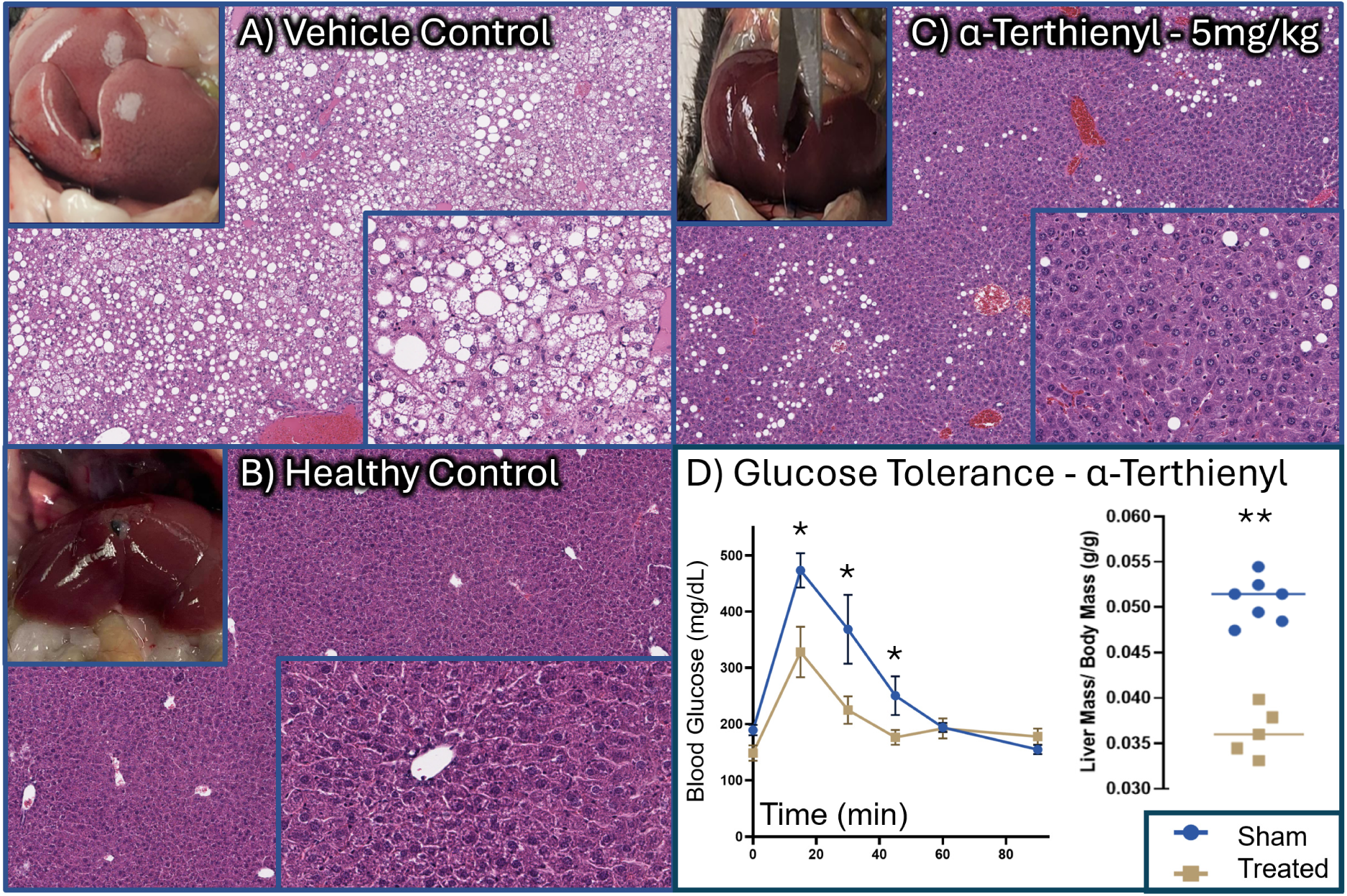
H&E-stained liver sections of A) Sham injection control, B) Healthy control, C) α-terthienyl administered for 12-days at 5mg/kg showing improvement in gross liver appearance (inset) and complete resolution of microvesicular steatosis with partial resolution of macrovesicular steatosis, D) improvement in glucose tolerance and decreased liver weight. (** denotes p<0.01)

Representative images of hematoxylin & eosin (H&E) stained liver sections for compound-treated, and vehicle-treated animals are included in **Figure 3**. Liver histology showed almost complete resolution of microvesicular steatosis and a significant reduction of macrovesicular steatosis relative to the vehicle control cohort. Food intake was monitored daily and there was no significant difference in chow consumption between vehicle and the treatment group. **Table 3** shows quantitative differences in measured parameters including body weight, liver mass, and histologic scoring of steatosis. Furthermore, α-terthienyl treatment significantly improved whole-body glucose tolerance in the mice, leading to lower blood glucose levels and a faster return to baseline following the glucose challenge compared to the control group.

**Table 3.** In vivo efficacy of α-terthienyl in DIO mice. Data represented as mean ± SEM (N=7 for vehicle, and N=5 for α-terthienyl treatment group, n.s. = not significant)

| Parameter | Vehicle | α-terthienyl - 5mg/kg | p-value |
| --- | --- | --- | --- |
| Body Weight (g) | 56.33 ± 7.37 | 50.29 ± 6.79 | 0.17 (n.s.) |
| Liver mass (g) | 2.93 ± 0.25 | 1.91 ± 0.21 | 2.20E-05 |
| Liver/Body Mass Ratio | 0.0578 ± 0.004 | 0.0392 ± 0.00459 | 7.30E-07 |
| Degree of Steatosis (%) | 85.8 ± 8.61 | 27.4 ± 9.55 | 6.14E-07 |

### 3.4 AI-Driven Target Deconvolution Predicts DPP-IV and HSD17β13 as Molecular Targets for α-Terthienyl

To elucidate the molecular mechanism underlying the prominent cellular and in vivo effects of α-terthienyl, we used the chemical structure of α-terthienyl and its phenotypic signature from the high-content screen as input to Plex to predict potential protein targets. Based on the saliency score and chain of evidence (**Figure 4**) dipeptidyl peptidase 4 (DPP-IV) and 17-β hydroxysteroid dehydrogenase 1 (HSD17β1) were the top-ranked targets. **Figure 4** shows selected targets from the chain of evidence (8 total targets identified) supporting the predicted association between α-terthienyl and its interaction with DPP-IV and HSD17β1. Plex initially identified HSD17β1, a member of the 17-beta hydroxysteroid family and is responsible for catalyzing the conversion of estrone (E1) into estradiol (E2) as a candidate target^65^ as it is structurally similar to known targets. However, based on literature, the liver specific HSD17β13 isoform has strong genetic links to MASLD protection. Specifically, in 2018, a link between HSD17β13 and MASLD was discovered through genome-wide association studies (GWAS), which showed a significant correlation between a loss-of-function (LoF) SNP rs72613567 and levels of ALT,^66,67^ and HSD17β13 is expressed in hepatocytes and is a lipid-droplet-associated protein implicated in lipid metabolism.^68,69^ We therefore focused on HSD17β13 as the more relevant target despite ∼20–25% sequence identity between the isoforms.

**Figure 4.**
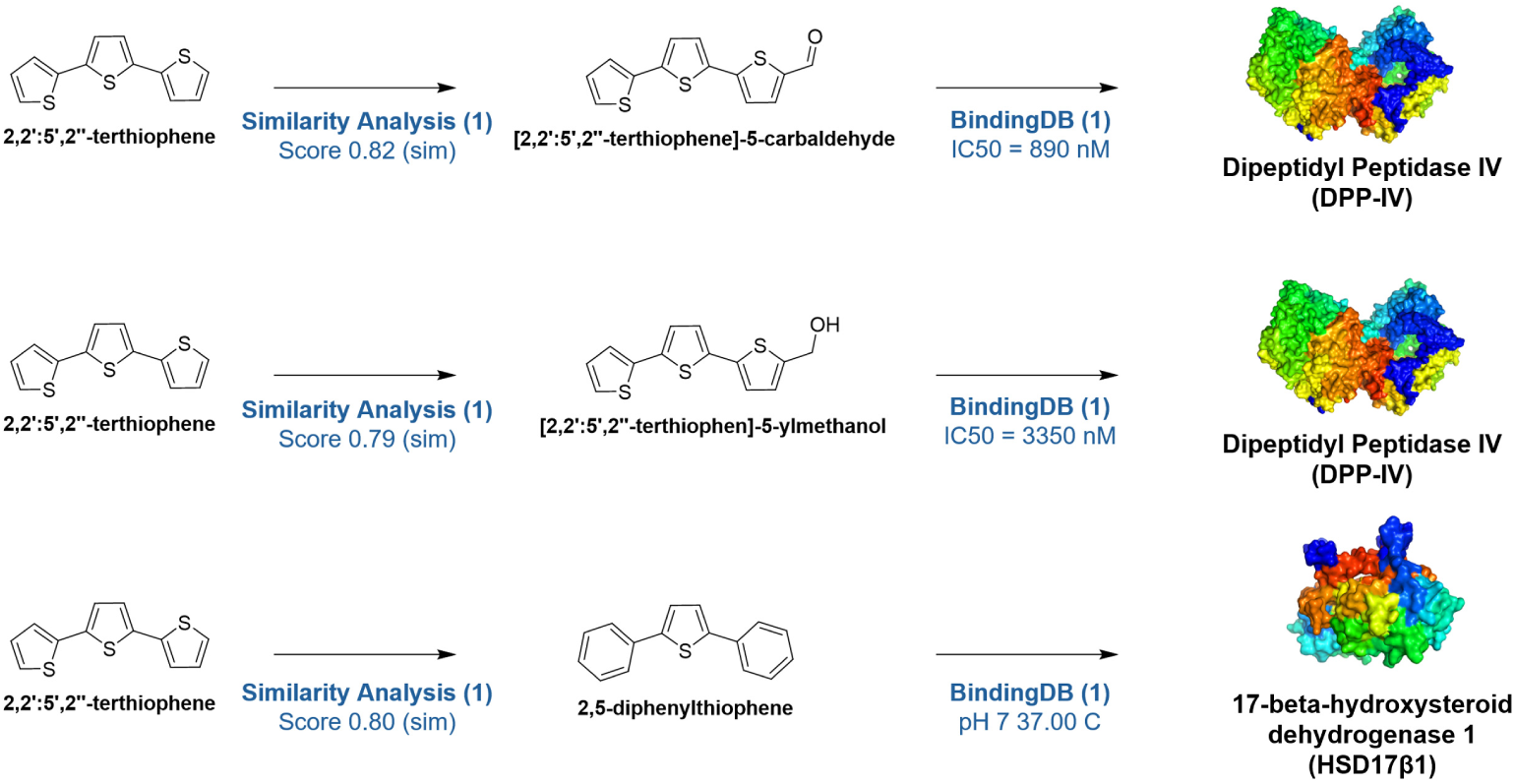
Chain of evidence showing selected targets linking α-terthienyl to DPP-IV and HSD17Β1.

### 3.5 Evaluation of DPP-IV and HSD17β13 as targets of α-Terthienyl

To evaluate the relevance of the targets predicted by AI we conducted deeper literature analysis into their signaling relevant to MASLD pathophysiology, molecular docking, and in vitro biochemical inhibition assays.

#### 3.5.1 Signaling Relevance of Dipeptidyl Peptidase 4 (DPP-IV) and GLP-1 Signaling for MASLD

DPP-IV is a widely expressed enzyme that rapidly inactivates incretin hormones, primarily glucagon-like peptide-1 (GLP-1) and glucose-dependent insulinotropic polypeptide (GIP).^70^ GLP-1, secreted from intestinal L-cells postprandially, plays a crucial role in glucose homeostasis by stimulating glucose-dependent insulin secretion from pancreatic β-cells, suppressing glucagon secretion from α-cells, delaying gastric emptying, and promoting satiety.^70^ Enhancing GLP-1 signaling, either through DPP-IV inhibitors (which prolong the action of endogenous GLP-1) or GLP-1 receptor agonists (GLP-1RAs), is an established therapeutic approach for T2DM and has shown promise in the treatment of MASLD/MASH.^71–73^ Beyond glycemic control, preclinical and clinical evidence suggests that enhancing GLP-1 signaling may also exert beneficial effects on the liver in the context of MASLD. Studies report that GLP-1RAs and DPP-IV inhibitors can reduce hepatic steatosis, decrease hepatic lipogenesis, increase fatty acid oxidation, and improve hepatic insulin sensitivity.^70^ The GLP-1 receptor is expressed on hepatocytes, suggesting potential direct liver effects.^70^ Furthermore, hepatocyte-derived DPP-IV itself may locally regulate portal GLP-1 bioactivity and influence hepatic glucose production.^74^ Targeting the DPP-IV/GLP-1 axis is therefore relevant for addressing the metabolic disturbances, particularly insulin resistance and hyperglycemia, that drive MASLD.

#### 3.5.2 Signaling relevance of Hydroxysteroid 17-beta Dehydrogenase 13 (HSD17β13) for MASLD

HSD17β13 is an enzyme predominantly expressed in hepatocytes and localized to lipid droplets.^75,76^ Its expression is often upregulated in MASLD.^77^ Genome-wide association studies (GWAS) have robustly linked loss-of-function variants in the *HSD17Β13* gene (notably the splice variant rs72613567:TA) with significant protection against progression to MASH, fibrosis, cirrhosis, and HCC in individuals with MASLD and other chronic liver diseases.^66,78^ This protective effect occurs despite these variants not typically altering simple steatosis levels, suggesting a role in downstream injury and fibrosis pathways.^79^ While its precise function is still under investigation, HSD17β13 has been proposed to act as a retinol dehydrogenase and is implicated in regulating hepatic lipid and phospholipid metabolism, potentially influencing lipid droplet dynamics and lipotoxicity.^76,80^ Some studies also link its activity or related phase separation phenomena to inflammation and pyrimidine catabolism pathways relevant to fibrosis.^79^ Consequently, inhibiting HSD17β13 activity is considered a promising therapeutic strategy for MASH and liver fibrosis.^75^

#### 3.5.3 α-Terthienyl Biochemically Inhibits Activity of DPP-IV and HSD17β13

To test if α-terthienyl can inhibit the enzymatic activity of these targets, we performed biochemical assays to assess α-terthienyl’s effect on DPP-IV and HSD17β13 activity. For DPP-IV, we quantified the conversion of the proluminescent Gly-Pro-aminoluciferin to luminescent aminoluciferin using purified enzyme using the DPP-IV-Glo™ Protease Assay kit. Diprotin A, a well-characterized DPP-IV inhibitor, was used as a positive control. We tested 9 doses in duplicate of α-terthienyl ranging from 100 µM–20 nM and found a clear dose-responsive relationship with an IC_50_ of 1.47 µM (**Figure 5A**). These results indicate that α-terthienyl acts as a direct inhibitor of DPP-IV.

**Figure 5.**
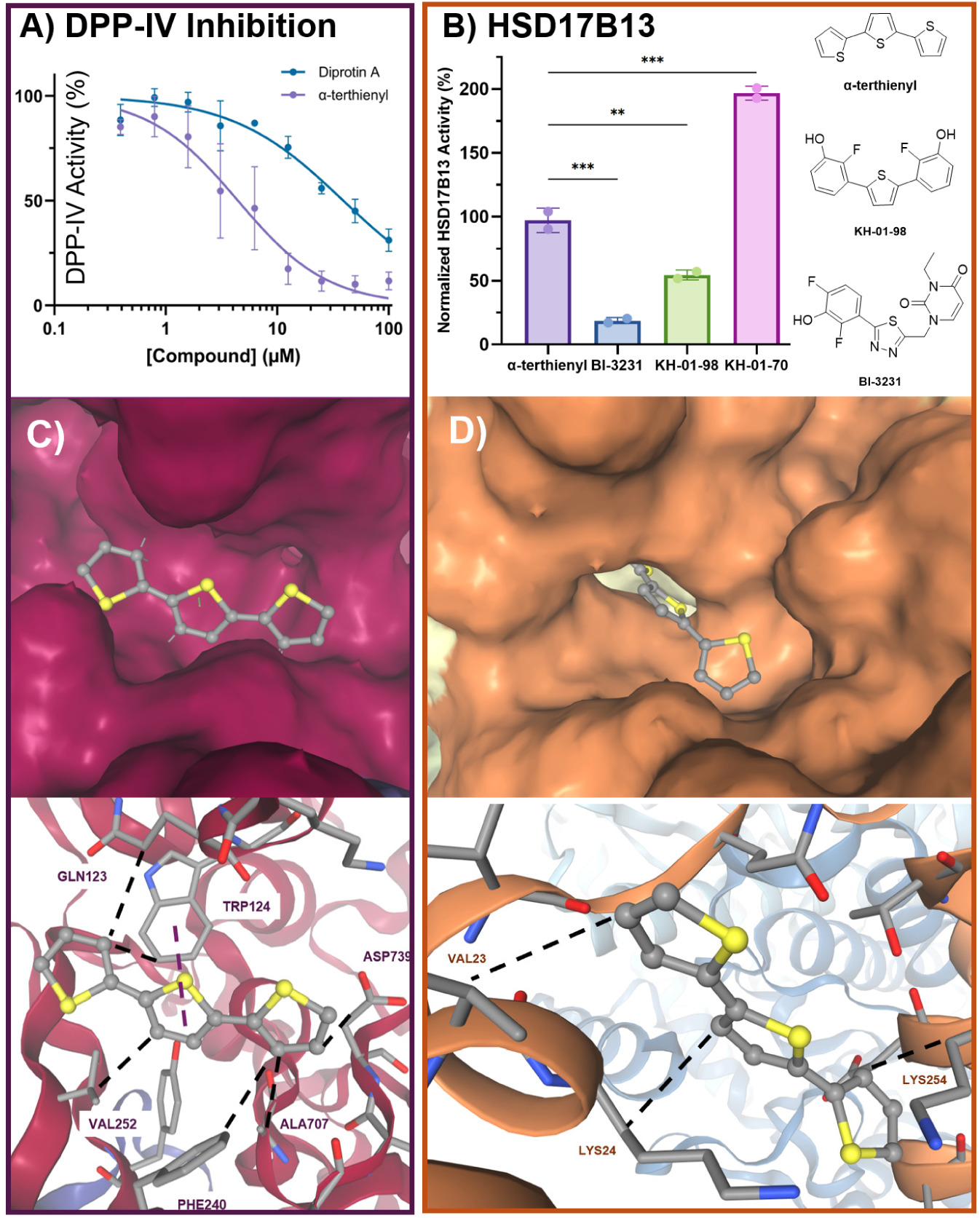
A) Biochemical inhibition of DPP-IV, B) In cellulo activity of HSD17β13, C) Docking of α-terthienyl into the DPP-IV binding pocket (PDB: 1NU6) using AutoDock Vina revealed a predicted binding affinity of –5.538 kcal/mol, with the compound participating in pi-stacking (purple) and hydrophobic (black) interactions situated in deep surface-accessible pocket. D) α-terthienyl docked into HSD17β13 (PDB: 8G9V), showing a favorable binding pose and predicted affinity of -5.686 kcal/mol, with hydrophobic interactions (black) deep within the surface-accessible pocket, consistent with its paradoxical enhancement of enzymatic activity.

Regulation of HSD17β13 activity, measured by estradiol turnover in cells (**Figure 5B**), revealed a nuanced interaction. While α-terthienyl itself showed minimal direct inhibition of HSD17β13 activity (97.2% activity relative to vehicle control), its oxidized/halogenated analog, KH-01-98 inhibited HSD17β13 activity to 54.4%. The known potent HSD17β13 inhibitor, BI-3231, served as a positive control and inhibited estradiol turnover to 18.7%, validating the cellular assay. These findings suggest that while α-terthienyl directly inhibits DPP-IV, its effects on HSD17β13 may be mediated through metabolic transformation or involve a distinct, yet related, structural motif.

### 3.6 Molecular Docking Supports Interaction of α-Terthienyl with DPP-IV and its Oxidized Metabolite with HSD17β13

To assess the structural feasibility of the AI predictions, molecular docking simulations were performed.^32^ Docking of the parent compound, α-terthienyl, into the active site of human DPP-IV revealed a plausible binding mode with favorable predicted binding energy (**Table 4**). The simulation showed potential interactions, such as hydrophobic contacts and possibly π-stacking, between α-terthienyl and key residues within the DPP-IV substrate-binding pocket, consistent with potential inhibition.

**Table 4.** Molecular docking results of α-terthienyl and an oxidized analog into DPP-IV (1NU6) and HSD17β13 (8G9V).

| Protein | Ligand | Predicted Binding Energy (kcal/mol) | Key Interacting Residues |
| --- | --- | --- | --- |
| DPP-IV | α-terthienyl | -5.538 | Trp124 (π-stacking), Val252, Gln123 |
| HSD17B13 | α-terthienyl | -5.686 | Lys24, Val23, Lys254 |
| DPP-IV | 2,5-diphenylthiophene 1,1-dioxide | -7.169 | Phe240 (π-stacking), Trp124, Gln123, Asp739 |
| HSD17B13 | 2,5-diphenylthiophene 1,1-dioxide | -6.664 | Gln29 (H bonding), Lys255, Arg276, Asn253, Ile280 |

Interestingly, docking α-terthienyl into the HSD17β13 active/substrate-binding site yielded less favorable scores or poses. However, considering that thiophenes can undergo metabolic oxidation, particularly S-oxidation, mediated by cytochrome P450 enzymes predominantly in the liver,^62^ we hypothesized that an oxidized metabolite might be the active ligand for HSD17β13. Docking simulations were therefore performed with an oxidized α-terthienyl analog. This analog, 2,5-diphenylthiophene-1,1-dioxide, showed a favorable predicted binding energy and formed plausible interactions within the HSD17β13 binding pocket, potentially interacting with residues near the catalytic site or cofactor binding region identified in structure-activity relationship studies.^81^ These differential docking results suggest a compelling hypothesis: α-terthienyl itself may primarily target DPP-IV, while its hepatic metabolite targets HSD17β13, providing a structural basis for the dual-target activity observed in vivo. Additionally, 20 small molecule decoys were docked into HSD17β13 and DPP-IV, and 2,5-diphenylthiophene-1,1-dioxide consistently exhibited strong binding affinity relative to the decoys (data not shown).

## 4. Discussion

In this study, we have integrated a high-content phenotypic screening with AI-driven target deconvolution to identify chemical modulators of steatosis and found a natural product of African marigold roots, α-terthienyl, as a novel potential therapeutic agent for MASLD. We began by screening the Selleck Chemicals Bioactive Compound Library-II containing 5,309 compounds for modulators of lipid accumulation in human hepatocytes. Using a ML score computed from cell-level morphological features, we identified 15 inhibitors and 16 exacerbators. Leveraging known targets for these drugs, clinical candidates, and probe compounds, we used the Plex AI platform to do a target enrichment analysis and identified biologically meaningful mechanisms related to hepatic steatosis. We determined that α-terthienyl had a potent in vitro IC_50_ ∼100 nM for steatosis inhibition and could ameliorate hepatic steatosis and improve glucose tolerance in a DIO mouse model of MASLD. Remarkably, daily IP administration of α-terthienyl at 5mg/kg body weight reduced the degree of steatosis from 85.8% (vehicle) to 27.4% (treatment) with nearly complete resolution of microvesicular steatosis. To explain potential mechanisms of action, we then used Plex to analyze molecular and literature-based evidence to predict targets, identifying DPP-IV and HSD17β13 as the top ranked candidates. To evaluate these predictions, we conducted a deeper literature evaluation, molecular docking, and biochemical testing. Together these analyses support DPP-IV and HSD17β13 as targets and future directions will focus on determining their causal role in MASLD.

To summarize the key elements of the potential roles for DPP-IV and HSD17β13 as targets of α-terthienyl mediating its effects on lipid accumulation: For DPP-IV as a target, GLP-1 agonists have been a breakthrough strategy to treat obesity and other diseases associated with metabolic syndrome and notably the successful clinical trial of semaglutide to treat NASH.^72^ DPP-IV is a key enzyme that degrades endogenous GLP-1, so its inhibition is expected to increase levels of active GLP-1.^82^ This aligns with the observed improvement in glucose tolerance in the DIO mouse model.^82^ Enhanced GLP-1 signaling improves glucose-dependent insulin secretion, suppresses glucagon, and can improve insulin sensitivity.^70^ Furthermore, direct effects of GLP-1 on hepatocytes, mediated via the GLP-1R, may contribute to reduced steatosis by decreasing lipogenesis and increasing fatty acid oxidation.^70^ Targeting DPP-IV addresses the systemic metabolic dysfunction, particularly impaired glucose regulation and insulin resistance, which is a defining feature and driver of MASLD.^83^ We found that α-terthienyl not only docks into DPP-IV but also has an in vitro IC_50_ of ∼1 uM. While this is 10-fold less potent than the phenotypic effects observed in our high-content screen, in the in vitro assay, the positive control Diprotin A, also has a 5–10x increase in IC_50_ compared to previous studies, suggesting that this might be an overestimate of the concentration for physiological relevance.

For HSD17β13 as a target, this HSD isoform has been genetically validated as a protective factor against MASLD/MASH progression and fibrosis.^84^ The mechanism by which HSD17β13 inhibition protects the liver may involve modulation of lipid droplet function, reduction of lipotoxic lipid species,^80^ alteration of retinol metabolism,^76^ or effects on inflammatory or fibrotic pathways, possibly via pyrimidine catabolism.^79^ Targeting HSD17β13 addresses aspects related to hepatic lipid handling and the progression towards more severe liver damage. While α-terthienyl did not dock in HSD17β13 and showed little inhibition of HSD17β13 in a cell based biochemical assay, we found that an oxidized metabolite of α-terthienyl did both. Thiophenes are known substrates for hepatic cytochrome P450 enzymes, often undergoing S-oxidation or epoxidation.^85,86^ This metabolic activation could potentially confer liver-targeted activity for HSD17β13 inhibition. The histologic pattern observed in vivo, characterized by a substantial resolution of microvesicular steatosis with persistent but diminished macrovesicular steatosis, is consistent with the expected biological consequences of HSD17β13 inhibition. HSD17β13 is a lipid-droplet–associated enzyme implicated in regulating hepatocellular lipid handling and lipotoxic injury rather than directly mediating bulk triglyceride storage.^76^ Genetic and pharmacologic studies suggest that loss of HSD17β13 function preferentially mitigates pathways linked to hepatocyte stress, aberrant lipid droplet remodeling, and the accumulation of small, dispersed lipid droplets associated with microvesicular steatosis.^76^ The treatment-associated clearance of microvesicular fat observed in our DIO mouse model aligns with this mechanistic framework, indicating improved hepatocellular lipid processing and reduced lipotoxicity. In contrast, macrovesicular steatosis, reflecting large, stable lipid droplets driven primarily by systemic metabolic inputs was only partially reduced, a pattern expected when intrinsic hepatocellular injury pathways are corrected in the absence of prolonged metabolic reprogramming. This selective histologic response supports a model in which HSD17β13 inhibition contributes to the restoration of hepatocyte homeostasis and attenuation of the more pathologic microvesicular phenotype while requiring longer or complementary metabolic interventions to fully resolve macrovesicular lipid accumulation.

While the separate engagement of DPP-IV (via parent drug) and HSD17β13 (via metabolites) by α-terthienyl represent potential mechanisms of action, the simultaneously targeting both may cooperatively treat MASLD. The parent compound and metabolites target two distinct, but highly relevant pathways implicated in the disease: one related to intrinsic hepatic processes and disease progression (HSD17β13) and the other related to the driving systemic metabolic disturbances (DPP-IV/GLP-1). Such multi-target polypharmacology may offer synergistic or additive benefits compared to selective inhibitors, potentially leading to greater efficacy in improving both liver health and metabolic control.^87^

The integration of PDD with AI represents a powerful strategy for modern drug discovery, particularly for complex diseases.^88^ PDD allows for unbiased identification of compounds active in relevant biological systems, capturing complex cellular responses that might be missed in target-based assays.^21^ However, the traditional bottleneck of identifying the MoA hinders the progression of phenotypic hits. AI-driven target deconvolution methods, like the Plex Research platform used here,^22^ can rapidly analyze phenotypic and chemical data to generate high-probability target hypotheses, significantly accelerating the path from hit identification to mechanistic understanding and preclinical validation. This study serves as an example of this synergy, where AI successfully predicted biologically relevant targets for a PDD hit, guiding subsequent validation efforts.

Several limitations of this study warrant acknowledgement. (1) pharmacokinetic (PK) data for α-terthienyl and its oxidized analog were not obtained. Understanding the absorption, distribution, metabolism, and excretion profile is crucial to confirm in vivo exposure, validate the proposed metabolic activation pathway leading to HSD17β13 inhibition, and assess potential liabilities such as P450-mediated drug-drug interactions or toxicity, which have been associated with some thiophene-containing drugs.^89^ (2) Direct evidence for the formation of the specific oxidized metabolite in vivo and its role in target engagement was not performed. (3) the efficacy was demonstrated in a single, albeit relevant, preclinical model (C57BL/6J DIO mouse); validation in other MASLD/MASH models with varying characteristics (e.g., more prominent fibrosis) would strengthen the findings. (4) While the in vitro potency for DPP-IV inhibition is consistent with the in vivo efficacy, it is ∼10 less potent than what was observed in the phenotypic screen that can be clarified in future metabolite-ID and PK studies. (5) While little toxicity was observed, a comprehensive off-target profiling of α-terthienyl and its metabolite was not performed that would enhance its potential for lead optimization.

## 5. Conclusion

In conclusion, this study successfully used an integrated drug discovery strategy, combining high-content phenotypic screening with AI-driven target deconvolution, to identify α-terthienyl as a novel anti-steatotic compound with promising activity against MASLD. In vivo studies confirmed its efficacy in ameliorating steatosis and improving glucose tolerance in a relevant mouse model of MASLD. AI predictions, supported by molecular docking, and biochemical confirmation suggest a dual-target mechanism involving inhibition of DPP-IV by the parent compound and inhibition of HSD17β13 by an oxidized analog, however, we attribute the majority of the improvement in liver histology to HSD17β13 inhibition. This polypharmacological profile, addressing both hepatic lipid handling and systemic metabolic control, positions α-terthienyl as a potential therapeutic lead for MASLD. The study highlights the power of integrating phenotypic screening with advanced AI tools to accelerate the discovery of therapeutics with complex mechanisms for multifactorial diseases. Further investigation into the pharmacology, mechanism, and safety of α-terthienyl is warranted.

## Acknowledgements

This work was supported by the National Institute of General Medical Sciences R01 award R01GM152417 (JZS), R01DK120623 (JZS and MCC), NIH OD (S10OD034245 to J.Z.S.), Pharmacological Sciences Training Program (PSTP) T32 training grant (JWW), NIH T32 (GM140223-02 to KMH).

## 6. Author Contributions

**Kathryn M. Hoegeman**: Methodology, Data curation, Writing. **Jesse W. Wotring**: Methodology, Data curation, Writing. **Reid Fursmidt**: Methodology, Data curation. **Jedidiah Gaetz**: Methodology, Data curation. **Ehab M. Khalil**: Software, Formal analysis, Data curation. **Douglas W. Selinger**: Software, Formal analysis, Data curation. **Tracey L. Schultz**: Methodology. **Sean M. McCarty**: Software, Validation, Data curation. **Matthew J. O’Meara**: Software, Validation, Data curation, Writing. **Martin C. Clasby**: Supervision, Project Administration, Review & Editing. **Jonathan Z. Sexton**: Conceptualization, Supervision, Formal analysis, Project administration, Writing – Review & Editing.

