## Supplementary Figures and Methods for "Phenotypic Screening Coupled with AI-Driven Target Deconvolution Identifies α-Terthienyl as a Dual DPP-IV/HSD17β13 Modulator with Efficacy in a Mouse Model of MASLD" (from another version of this preprint: version 1, 2025-12-16)

### Supplementary Data/Figures

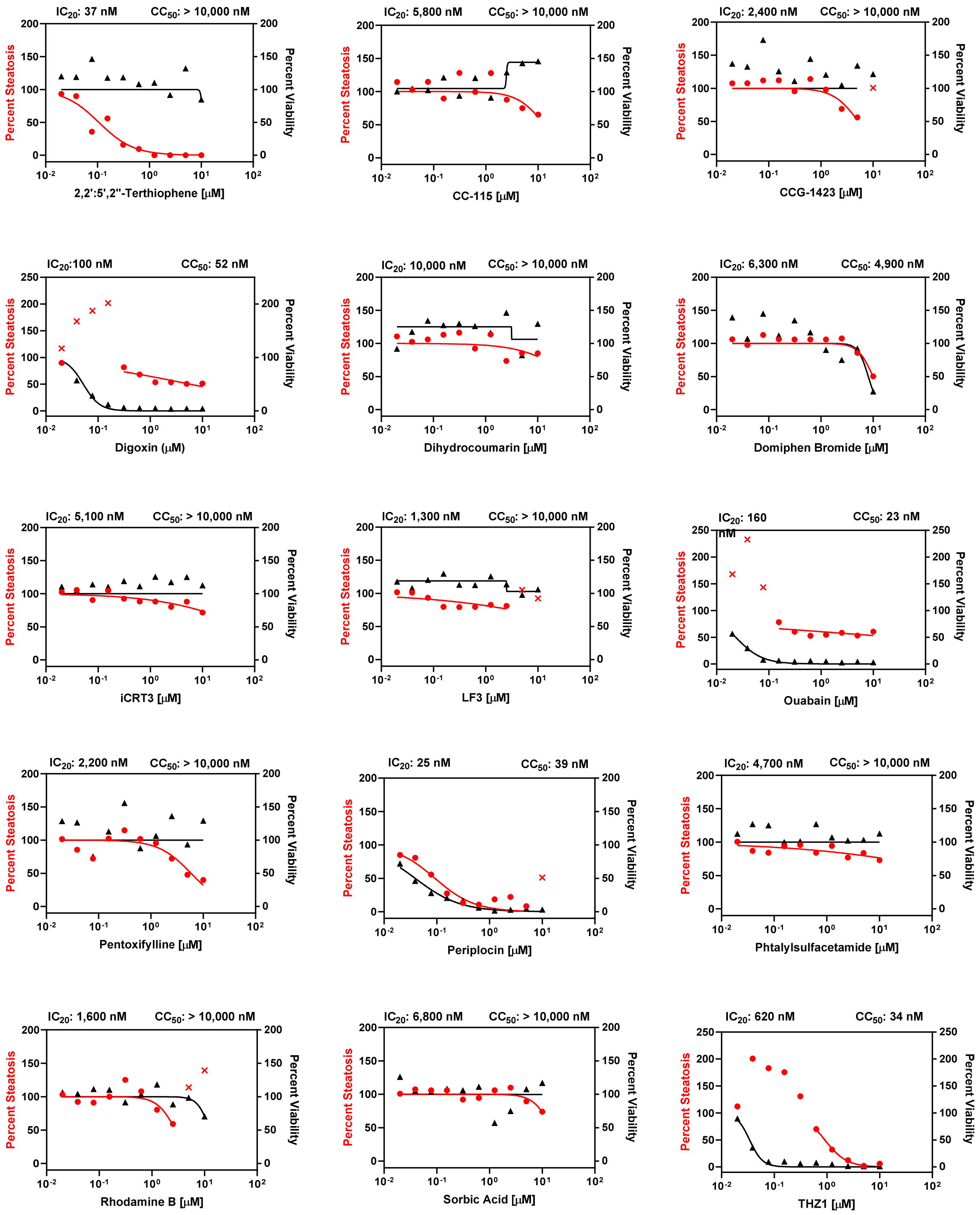


**Supplementary Figure 1**. Representative concentration–response curves for hit compounds that reduced intracellular lipid accumulation in the steatosis assay. For each compound, percent steatosis (red circles, left y-axis) and percent viability (black triangles, right y-axis) were quantified across a serial dilution range. $IC_{50}$ values for steatosis reduction and $CC_{50}$ values for cytotoxicity are reported above each panel. Compounds display a range of potencies and therapeutic windows.

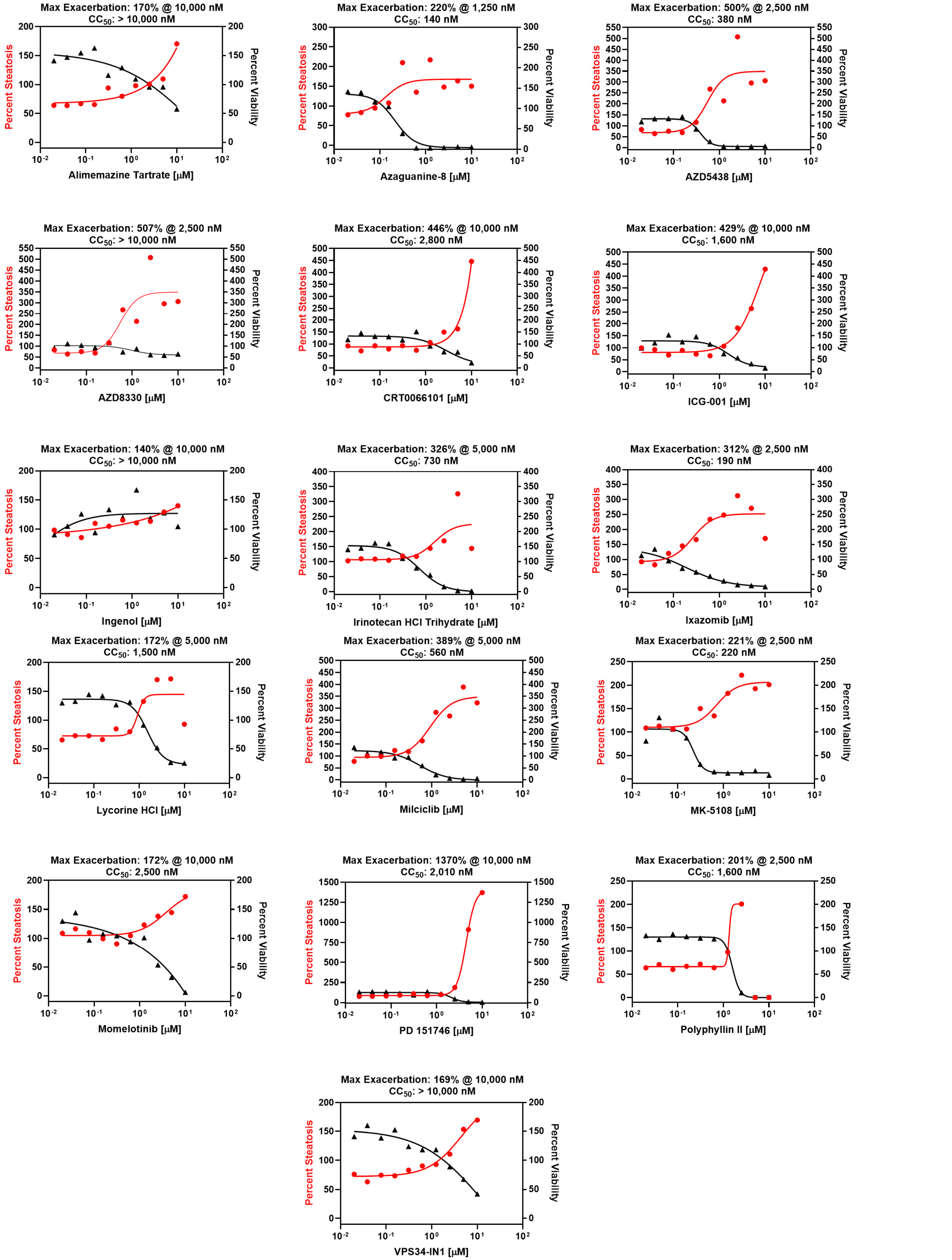


**Supplementary Figure 2**. Representative concentration–response curves for hit compounds that exacerbate intracellular lipid accumulation in the steatosis assay. For each compound, percent steatosis (red circles, left y-axis) and percent viability (black triangles, right y-axis) were quantified across a serial dilution range. $IC_{50}$ values for steatosis reduction and $CC_{50}$ values for cytotoxicity are reported above each panel. Compounds display a range of potencies and therapeutic windows.

**Supplemental Methods – Synthesis of KH-01-98**

**General Chemistry Methods.** All reagents were used as received by the manufacturer without further purification. We purchased α-terthienyl (CAS: 1081-34-1) from MedchemExpress. Reactions were followed by thin layer chromatography (TLC) performed on Silica gel 60G F254 glass plates. Visualization was performed using ultraviolet light at 254 nm. Flash chromatographic purifications were performed using a Teledyne ISCO Combiflash RF with Redisep Gold columns. An Agilent 1100 series HPLC with an Agilent Zorbax Eclipse Plus–C18 column was used to determine purity. All compounds were determined to be >95% pure by HPLC analysis with detection at 254 nm. Products were verified by $^{1}H$ and $^{13}C$ NMR in DMSO-$d_6$ at room temperature on Varian Inova 500 MHz instruments. Mass spectrometry data were collected on an Agilent 6320 Q-TOF HPLC-MS or Micromass AutoSpec Ultima mass spectrometer.

**Synthesis of Analogs KH-01-85 and KH-01-98**

***3,3'-(thiophene-2,5-diyl)bis(2-fluorophenol) (KH-01-98).*** The title compound was prepared following established Suzuki-coupling conditions[1] using 2,5-dibromothiophene (121 mg, 0.500 mmol), 2-fluoro-3-hydroxyphenyl)boronic acid (312 mg, 2.00 mmol), and tetrakis(triphenylphosphine)-palladium(0) (28.9 mg, 25 μmol). Reagents were combined in a 25-mL round bottom flask fitted with a stir bar and reflux condenser. The system was purged with nitrogen for 10 min. A purged solution of aqueous sodium carbonate (0.1 g/mL, 3 mL) was then added, and the reaction mixture was refluxed overnight with vigorous stirring under nitrogen. After cooling, the reaction was diluted with water and extracted with diethyl ether (3 x 10 mL). Combined organic layers were washed sequentially with water (10 mL) and brine (10 mL), dried over sodium sulfate, and evaporated to dryness. The crude product was purified by column chromatography using hexanes to afford the desired compound, yielding the desired product as a yellow solid (27 mg, 18%). $^{1}H$ NMR (500 MHz, DMSO) δ 10.06 (s, 2H), 7.57 (s, 2H), 7.19 (t, J = 7.2 Hz, 2H), 7.05 (t, J = 7.9 Hz, 2H), 6.92 (t, J = 8.2 Hz, 2H). $^{13}C$ NMR (126 MHz, DMSO) δ 147.18, 146.30 (d, J = 12.4 Hz), 137.13, 127.45 (d, J = 5.2 Hz), 125.20 (d, J = 4.8 Hz), 122.30 (d, J = 10.0 Hz), 118.28, 117.38. ESI-TOF *m/z* [MH]- calcd for $C_{16}H_{10}F_2O_2S$ 304.03696 found 303.0293

***2,5-diphenylthiophene-1,1-dioxide (KH-01-85).*** A solution m-CPBA (59 mg, 0.34 mmol, 2.7 eq.) in DCM (0.7 mL) was added dropwise to 2,5-diphenylthiophene (30 mg, 0.13 mmol, 1 eq.) in DCM (0.7 mL). The reaction mixture stirred at room temperature for 72 hours. Upon reaction completion, the crude mixture was diluted with DCM and washed with water (3x10 mL) and brine (10 mL) then dried over sodium sulfate. The organic material layers were combined, reduced to dryness, and purified via column chromatography using a 24 g silica gel column with gradient of 0% to 5% MeOH/DCM to afford 2,5-diphenylthiophene-1,1-dioxide as a yellow solid (13 mg, 37%). $^{1}H$ NMR (500 MHz, DMSO) δ 7.83 (d, 4H), 7.67 (s, 2H), 7.53 (dt, J = 26.9, 7.4 Hz, 6H). $^{13}C$ NMR (126 MHz, DMSO) δ 140.48, 130.68, 129.87, 127.36, 126.50, 123.19. EI-MS m/z [M] calcd for C16H12O2S 268.05580, found 268.0558.

### NMR Spectra

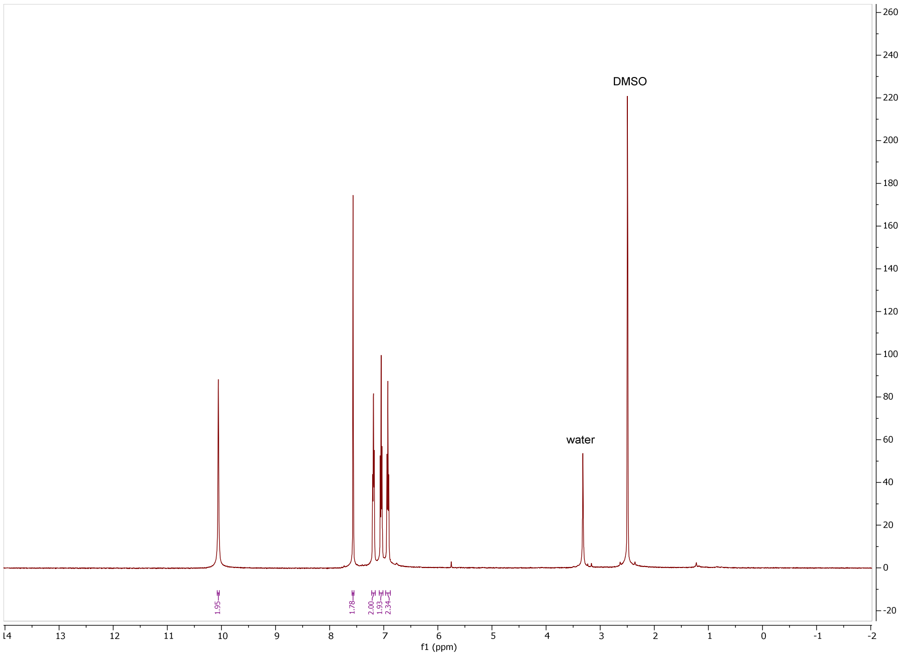


**Spectrum 1.1** $^{1}H$ NMR of KH-01-98 in DMSO.

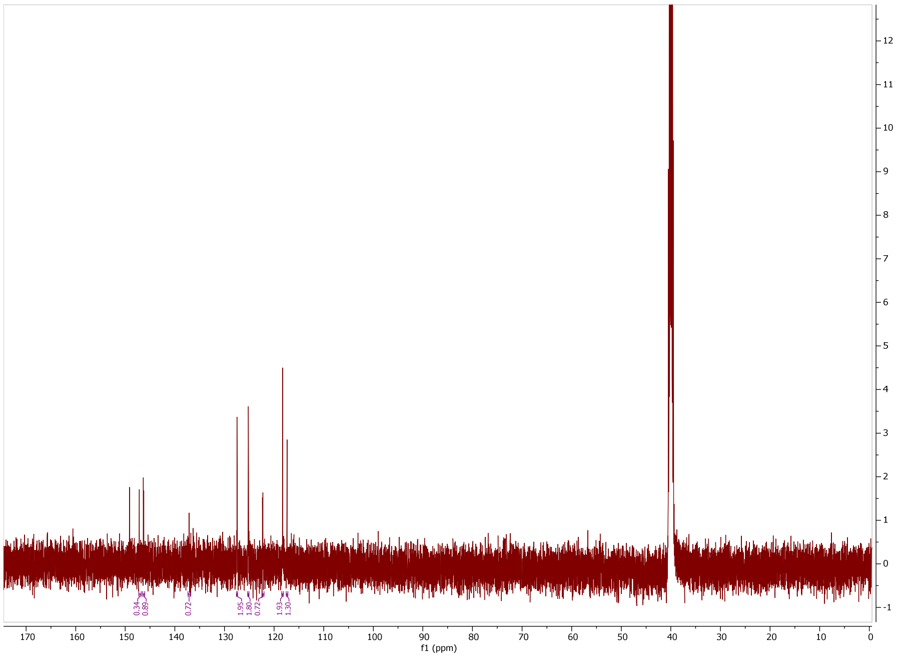


**Spectrum 1.2** $^{13}C$ NMR of KH-01-98 in DMSO.

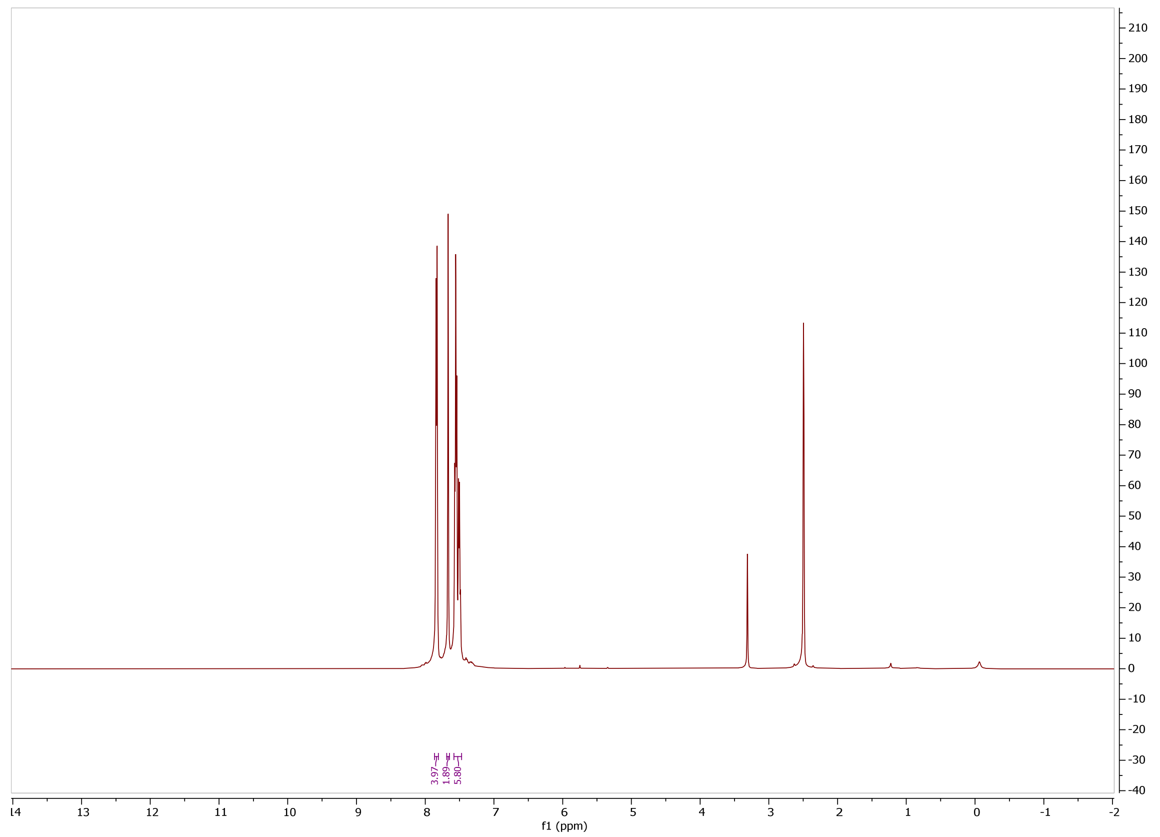


**Spectrum 1.3.** $^{1}$H NMR of KH-01-85 in DMSO.

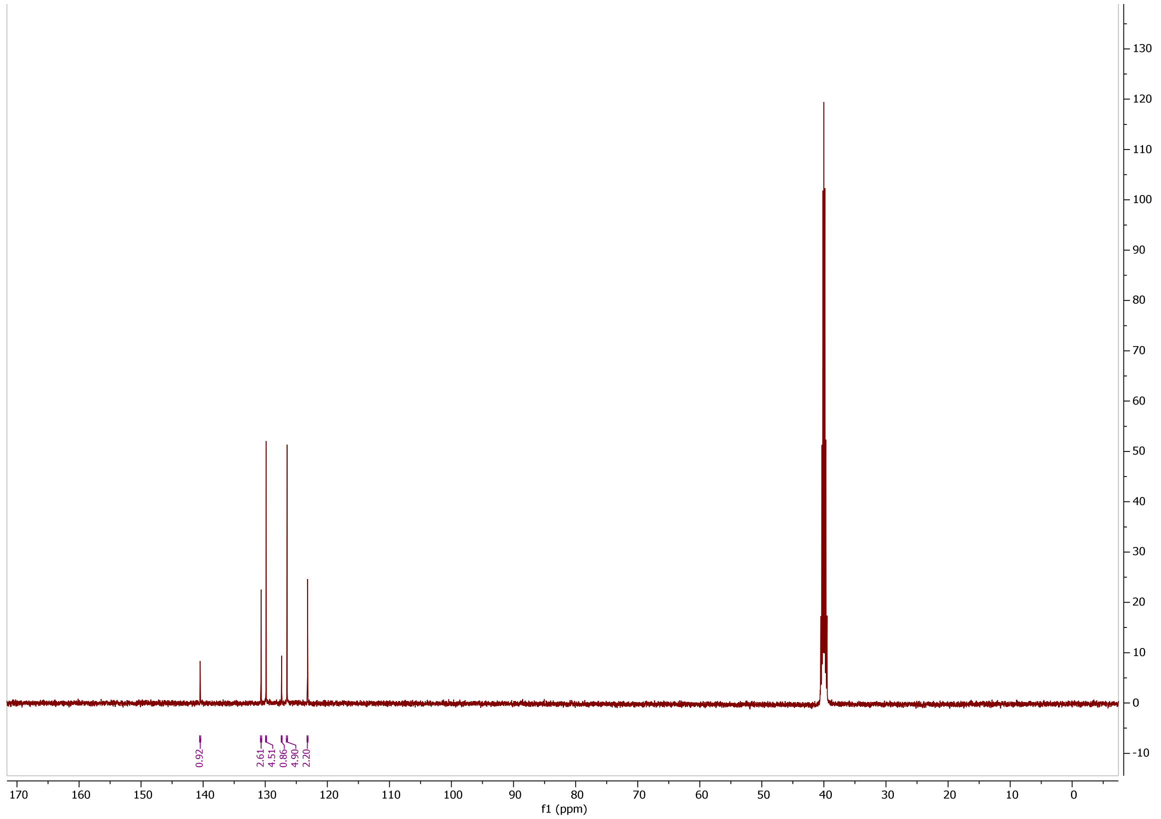


**Spectrum 1.4.** $^{13}$C NMR of KH-01-85 in DMSO.
